# Optimal harvesting strategies for ecological population dynamics

**DOI:** 10.1101/2023.04.04.535628

**Authors:** Sayeh Rezaee, Cesar Nieto, Abhyudai Singh

**Author notes:** {, }. { }.

## Abstract

What is the optimal way to harvest an ecological population sustainably is a fundamental problem in natural resource management. Here we use the framework of the stochastic logistic model which captures random birth-death of individuals to determine the optimal harvesting strategy that maximizes the integrated yield over time. Harvesting is assumed to occur at either a constant or state-dependent rate, and individuals are harvested with a certain probability whenever a harvesting event occurs. A special case of state-dependent harvesting is a threshold-based strategy, where harvesting is done when the population crosses a threshold. We use moment closure schemes to develop analytical formulas quantifying the mean and optimal yield. Moreover, as populations are susceptible to extinction at high harvesting rates, the Finite State Projection (FSP) method is used to estimate the probabilities of extinction across strategies and parameter regimes. Our results show that the threshold-based strategy is most effective in maximizing the yield as it suppresses population fluctuations and minimizes extinction events.

## I. Introduction

The problem of determining the optimal time to harvest a population that is growing dynamically is a common challenge in ecology and resource management [1]–[3]. To obtain an effective harvesting strategy, it is crucial to consider not only the economic benefits of the harvest but also the ecological sustainability of the population being harvested [4]. Over-harvesting can result in natural resource depletion, environmental degradation, and a loss of biodiversity [5]. In contrast, under-harvesting can lead to missed economic opportunities and reduced food security [6].

Modeling the population dynamics has been part of the challenge in the current approach to the problem. Previous contributions to the literature have focused on analyzing the optimal harvesting strategies for populations with deterministic continuous growth [7], [8]. Other approaches have considered stochasticity in population dynamics as a diffusive process [9]–[13]. However, these approaches have not considered the possibility of population extinction since they take the population as a continuous variable that can take any arbitrary small amount and almost never reaches zero [14]. As a result, such models have a limited approach to the problem of extinction from an ecological standpoint [15]. The harvesting process is an additional challenge in the problem description. Some approaches consider harvesting as a perturbative process where the population can be approximated as continuous [9], [10]. A more accurate model should take harvesting as a non-continuous jump where a large amount of resources can be collected [16], [17]. The amount of harvested resource is also subject to random variability since each individual has some probability of not being caught. Therefore, there is a need for more sophisticated models that can consider the complex dynamics of population growth, harvesting, and extinction risk, which can have significant implications for conservation and sustainable resource management.

This article studies different harvesting strategies for an integer-valued population with demographic stochasticity. For modeling the population dynamics, we use the discrete logistic growth model with intrinsic random fluctuations given by stochastic birth and death events [18]. While the rate of birth is proportional to the population count, the death rate is a quadratic function of the population. In the stochastic model, extinction is a certain outcome. However, the time to extinction can be significantly long for many specific parameter regimes [15], [19].

The harvesting process is modeled as a jump process where each individual has a certain probability of being caught. Different strategies can be distinguished by the frequency of the harvest. In this study, we focus on three strategies. The constant Poisson rate technique involves harvesting as a stochastic process with Poisson arrivals. The state-dependent rate strategy takes the probability of harvesting over a time interval increasing with the population count, while the threshold-based strategy triggers harvesting only once the population reaches a critical level.

The constant harvesting rate strategy is revisited from [20] providing analytical approximations to the maximum total yield and the optimum harvest rate. For state-dependent harvesting, we derive formulas for the optimal rate and yield using the moment dynamics approach, as previously described in the literature [21]–[25], by applying closure schemes to simplify the differential equations of the statistical moments [26]–[34]. For threshold-based harvesting, due to singularities in the moments dynamics, we use the time-averaging method. We should note that in our analytical approaches, we assumed a negligible probability of population extinction. To assess the validity of these analytical approximations, we numerically solved numerically the system while considering a non-zero probability of extinction. To do this, we used the finite state projection (FSP) method [35].

The article is structured as follows: In Section II, we present the problem formally and discuss the results for a constant harvesting rate. In Section III, we explore the state-dependent harvesting strategy and how to maximize the yield by optimizing the harvesting rate. Next, in Section IV, we explain the threshold-based strategy and optimize the threshold for maximizing the yield. Finally, we compare the different strategies and show that the threshold-based strategy is the most effective as it minimizes population fluctuations and extinction events, leading to maximum yield.

## II. Stochastic model formulation of harvesting

Let the integer-valued random process *x*(*t*) ∈ {0, 1, 2, … } represent the population count of the harvested species at time *t*. As explained in the schematic model in Fig. 1, our model considers the population count to evolve as a random birth-death process. The overall stochastic dynamics of *x*(*t*) is represented by the following resets that occur at random times, and when the reset occurs the corresponding rest map is activated

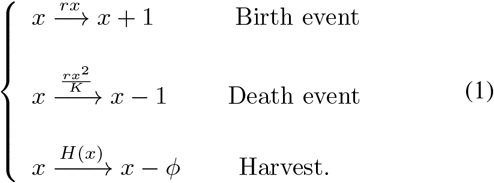

The first reset in (1) is the birth of an individual that occurs probabilistically with rate *rx*, where *r* is the exponential growth rate. Similarly, the death of an individual (a transition from *x* → *x* −1) occurs at a rate *rx*^2^*/K* with *K* being the carrying capacity. Finally, the last event corresponds to harvesting occurring with a state-dependent rate *H*(*x*). During each harvest, *ϕ* individuals are removed from the population. If each individual has a probability *f* of being harvested, then *ϕ* is a binomially-distributed random variable with parameters *x* and *f*. These individuals are added to the yield *y*(*t*) that is a positive real-valued random process. Between two consecutive harvesting events, the yield is assumed to decay as per first-order kinetics that captures its consumption

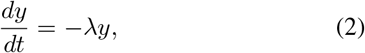

where *λ* is the consumption rate. Additionally the harvest reset map can be shown as

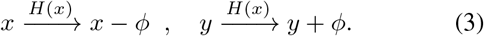

Note that in the absence of harvesting and assuming no population extinction, demographic stochasticity arising from random birth-death will cause *x*(*t*) to fluctuate around its carrying capacity.

**Fig. 1:**
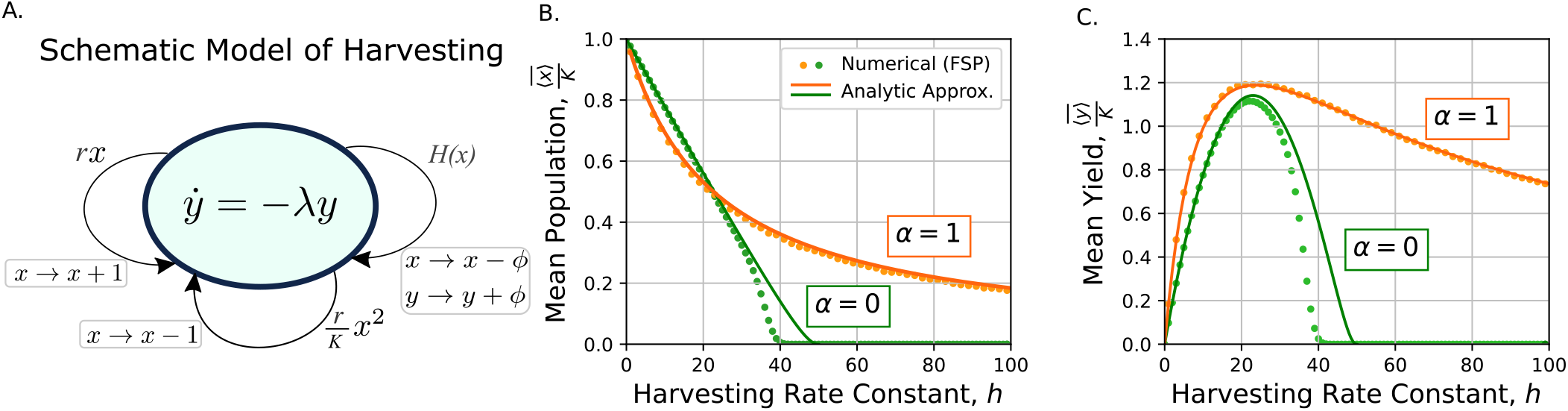
Harvesting at a state-dependent rate (*α* = 1) leads to a higher yield than harvesting at a constant rate (*α* = 0). **A**. In the proposed model, population *x* follows a logistic stochastic discrete dynamics. The yield *y* decreases exponentially over time due to consumption at an exponential rate *λ*. Each harvest occurs at a state-dependent rate of *H*(*x*) = *h* (*x/x*_*h*_)^*α*^ and results in a decrease/increase in population/yield by a random variable *ϕ*. **B**. Normalized mean population 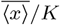 as a function of the harvesting rate constant *h*. **C**. Normalized mean yield 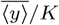 as a function of harvesting rate constant *h*. The results are represented for both constant Poisson harvesting rate (*α* = 0, green) and state-dependent rate (*α* = 1, orange). Analytic approximations correspond to the analytical results explained in Section III, while numerical results are obtained by numerically solving the system using the finite-state projection (FSP) algorithm for *tmax* = 400/*r* and *I* = 1200. The values of 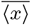 and 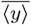 are normalized to the carrying capacity *K*. The rest of the parameters are taken as *r* = 10, *K* = 10^3^, *x*_*h*_ = 0.5 × 10^3^, *f* = 0.2, *λ* = 2.

In (1) and (2), the particular definition of the harvesting rate *H*(*x*) defines different harvesting strategies. In this article, we adopt the power-law function

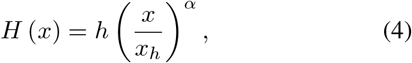

that is defined by three parameters - the exponent *α* ≥ 0, harvesting rate constant *h* > 0, and the harvesting threshold 0 < *x*_*h*_ < *K*. We will investigate three particular cases:

- A constant harvesting rate (*α* = 0).
- A harvesting rate that linearly increases with population size (*α* = 1).
- A threshold-based strategy of harvesting every time the population exceeds the threshold *x*_*h*_ (*α* → ∞) [36].

Ignoring stochasticity, in the simple deterministic limit of population counts evolving as per an ordinary differential equation, the maximum yield 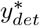 and its corresponding population density 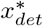 are as follows (see [20] for details)

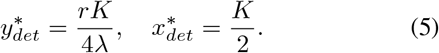

However, in practice, the population is subject to perturbations, some intrinsic, coming from individual birth and death, and others extrinsic, from the harvesting events. To determine the optimal yields in the stochastic formulation we first start by ignoring the effects of extinction by assuming a large population size. Using the Dynkin’s formula [37] for the Stochastic Hybrid System defined by (1)-(2), the mean yield ⟨*y*⟩ evolves as

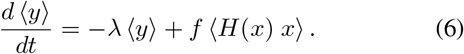

Throughout this manuscript we use angular notation ⟨ ⟩ for taking averages of random variables and processes. From (6), and assuming no population extinction, the steady-state mean yield 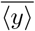 is given by

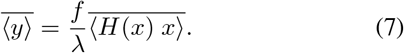

In the following sections, we investigate the population statistics and optimal yield for different harvesting strategies. We start with the the case of constant harvesting rate (*α* = 0).

### A. Constant harvesting rate (α = 0)

In this case, the harvesting rate is constant *H*(*x*) = *h*. This means that the harvesting events occur according to a Poisson process with a rate *h*. In the previous work [20], we presented moment dynamics for this case using Dynkin’s formula [37].

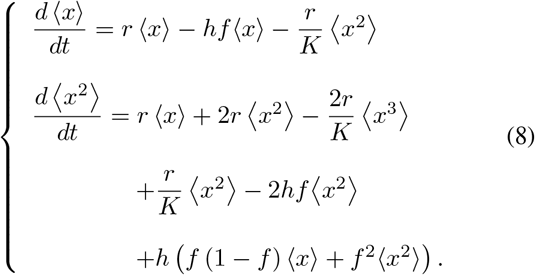

The high-order moments on the right-hand side of the above equation have led to an unclosed system of moment dynamics. This is because the dynamics of second-order moment ⟨*x*^2^⟩ depends on third-order moment ⟨*x*^3^⟩. To overcome this issue, the derivative matching moment closure scheme was proposed in a previous study [38], which approximates

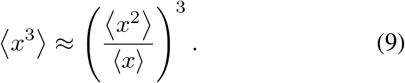

Then the steady-state population statistics can be approximated as the following formulas.

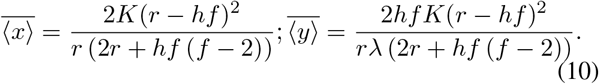

Figs. 1B and 1C show the trends of 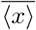 and 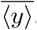, respectively, for a fixed *f* and an increasing *h*. The significance of extinction becomes evident when comparing the results in (10) with the numerical solution obtained using the finite state projection algorithm (will be discussed in the next section).

The optimal harvesting rate *h*\* and the corresponding maximum yield 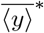 can be achieved as follows.

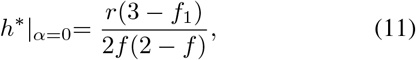

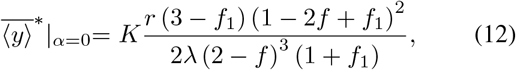

where 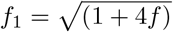 (see [20] for details).

Observe how, at the limit of the very low harvest probability *f* → 0, the optimal harvest frequency

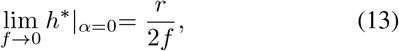

diverges by maintaining *fh*\* = *r/*2. In this limit, the optimal populations 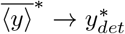 and 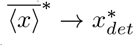 converge to their deterministic counterparts (5).

## III. state-dependent harvesting rate (*α* = 1)

This section presents the solution for the simplest state-dependent harvesting strategy *α* = 1 in (4). Therefore, the harvest rate can be written as 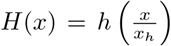. Following this strategy, harvesting is a random process with a rate that grows with the population.

### A. Analytical Approximation

To find the population statistics, we write the moment dynamics using Dynkin’s formula.

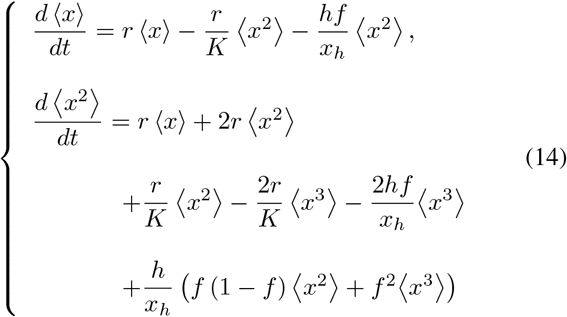

We use moment closure approximation (9) to get a closed system. On the other hand, for analytical tractability, we approximate the statistics at the limit of large population sizes, which leads to ⟨*x*^2^⟩ ≫ ⟨*x*⟩. Then, we can use the approximation below to simplify the moment dynamics in (14).

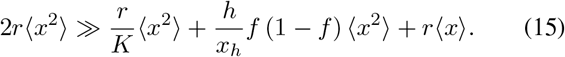

Consequently, the following steady-state moments for *x* are achieved.

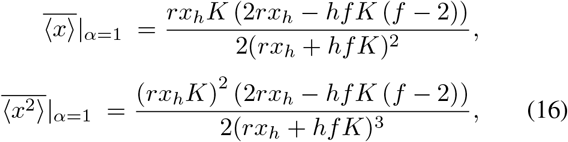

which give the steady-state mean yield from (7).

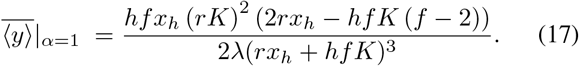

Fig. 1 shows the comparison between the mean yield obtained with the constant harvest rate (10) and the state-dependent rate (17). Observe how the mean yield is also a concave function of *h*. Hence, given *f*, it is possible to find the optimal harvesting rate *h*\* and the corresponding maximum mean yield 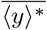 in steady state is calculated.

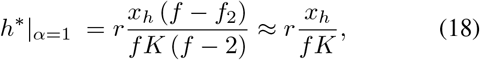

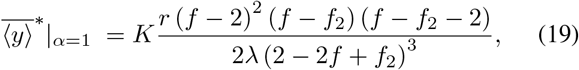

where 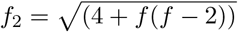.

Two main findings can be highlighted. First, as Fig. 1C shows, the expected yield for *α* = 1 is always greater than the one for *α* = 0 with the same parameters. Second, at the limit of perturbative harvest (*f* → 0)

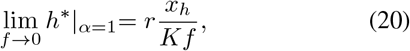

which diverges similarly as (11). This time, the optimum harvesting frequency scales with *f* such as *fh*\* = *rx*_*h*_*/K*. With this optimal rate, 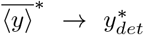 and 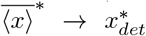 converge to their deterministic counterparts (5). Next, we will solve the system numerically using the Finite-State Projection algorithm to estimate the validity of the analytical approximations.

### B. Numerical Solution

The birth-death process described in (1) can be modeled using the forward Chapman-Kolmogorov formulation [39], also known as the Master equation. In this framework, the probability vector *P* (*t*) with components *p*_*i*_, *i* ∈ {0, 1, 2, … } represents the probability of existing *i* individuals at time *t*. To approximate a numerical solution, we propose to use the Finite State Projection (FSP) algorithm [35], which truncates infinite number of equations in the Master equation. Considering the first *I* elements of *P* (*t*) = [ *p*_0_(*t*) *p*_1_(*t*) … *p*_*i*_(*t*) … *p*_*I*_ (*t*) ]^*T*^ gives the Master equation as follows.

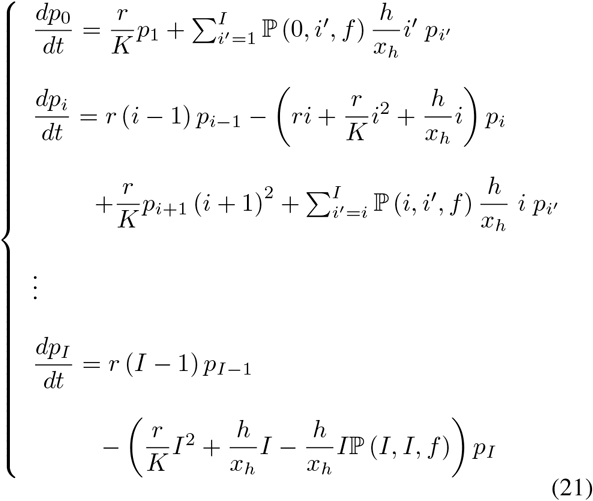

ℙ (*m, n, f*) in the above equations represents the probability mass function of the Binomial distribution, i.e., the probability of obtaining a specific number of successes *m*, in a series of independent Bernoulli trials *n*, where the probability of success in each trial is denoted as *f*.

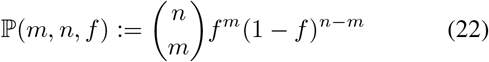

From (21), the dynamics of *P* (*t*) can be written as a linear system

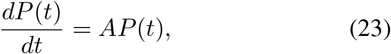

where *A* is an (*I* + 1) dimensional matrix. Consequently, *P* (*t*) is achieved as

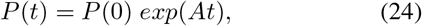

where *P* (0) = [0 0 … 1 … 0 0] is the vector of probabilities at time *t* = 0. We assume that the population starts in the state *x* = *K*. Now, the *n* − *th* order moment of *x* denoted by ⟨*x*^*n*^⟩ can be calculated from 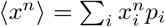.

Fig. 1 shows the comparison between the statistics estimated from analytical formulas (10) and (17) along with the optimal value of *h*\* and 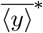. We also represent the results of the FSP to investigate the effects of extinction. It can be seen in Fig. 1B and 1C that analytical estimations provide an accurate approximation of the model with extinction for the given parameters during the studied time-scale (*t*_*max*_ < 400*/r*), where *t*_*max*_ is the time at which the statistics were calculated. The deviations between the numerical result and the analytical approximation can be attributed to extinction. The reason behind the fact that the state-dependent rate is more accurate is that in this case harvests occur rarely at small population numbers, which decreases the probability of extinction. In the next section, we consider the threshold-based harvest strategy, where the probability of harvest depends strongly on the population level.

## IV. Threshold-based strategy (*α* → ∞)

As shown in Fig. 2A, in the threshold-based strategy, the harvest occurs whenever the population exceeds the threshold population *x*_*h*_. During each harvest, the population decreases by the amount of *ϕ*, drawn from a binomial distribution with parameters *x* and *f*.

**Fig. 2:**
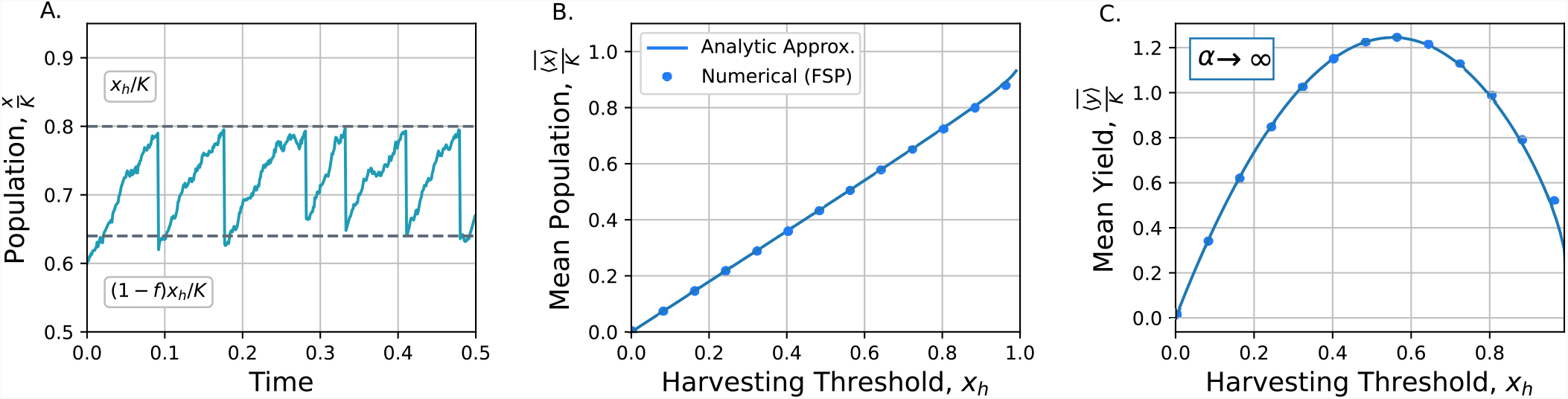
The mean yield for threshold-based harvest strategy (*α* → ∞) can be optimized over the threshold *x*_*h*_. **A**. An example of normalized population trajectory over time for *f* = 0.2, *x*_*h*_ = 0.8 *K*. **B**. Normalized mean population 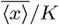 as a function of threshold *x*_*h*_. **C**. Normalized mean yield 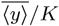 as a function of the harvesting rate constant *x*_*h*_. Values are normalized over the carrying capacity *K*. In the simulation, we considered the time *t* in FSP as *t*_*max*_ = 400*/r*. Other parameters are taken as *r* = 10, *K* = 10^3^, *λ* = 2.

### A. Analytical Approximation

As we discussed earlier, this case is equivalent to considering the limit value of *α* → ∞ in the general harvesting rate defined in (4). The moment dynamics approach we used to solve the system in the two previous cases is not applicable here due to the singularities in (4) at this limit. Therefore, we estimate the moments of the population taking the average over the time for a single trajectory. The steady-state mean population follows (see the Appendix for details and the explicit expression).

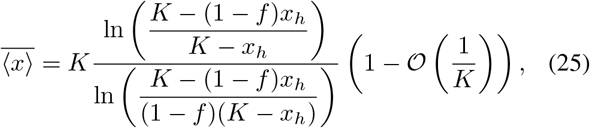

where the effect of randomness on the capture of each individual is included in 𝒪(1*/K*), with 𝒪 (*x*) standing for *order of* notation. In the limit of large *K*, we can neglect this term as 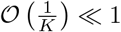. Finally, the mean yield of the stochastic system with threshold-based harvesting rate is computed as:

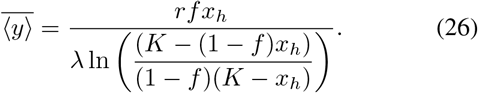

The population statistics for threshold-based harvesting as a function of threshold *x*_*h*_ is presented in Fig. 2B and 2C. Observe how 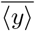 is a convex function of *x*_*h*_. Hence, given *f*, we can obtain numerically the value of 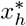 that maximizes the mean total yield in steady state 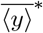 (see Fig. 2).

In the limit of perturbative harvest (*f* → 0), 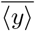 can be approximated as:

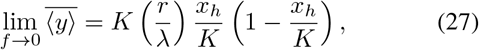

which is maximized for 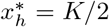. In this limit, similar to the other harvesting strategies, populations 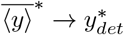 and 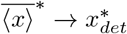 converge to their deterministic counterparts (5). To investigate how important the extinction is in this strategy, we will solve the system using the FSP algorithm.

### B. Numerical Solution

To analyze the exact behavior of the system considering the probability of extinction, we use the FSP method. Accordingly, we write the chemical Master equations for the stochastic harvesting model with threshold-based rate.

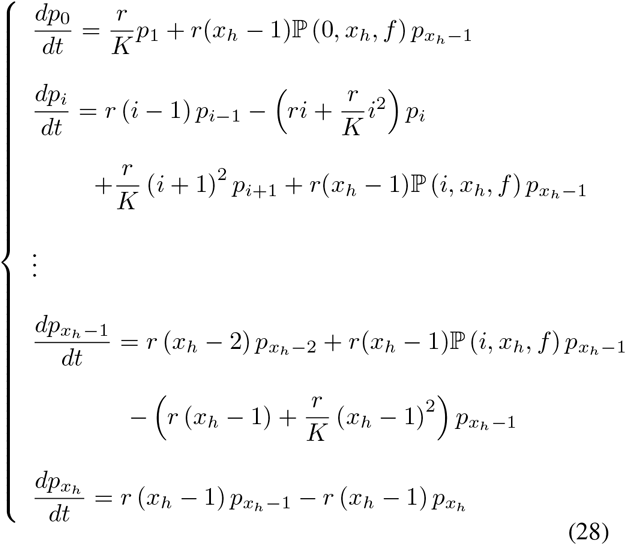

where 1 ≤ *i* ≤ *x*_*h*_, and *x*_*h*_ is the number of individuals at which the harvesting takes place. By solving the above set of equations, as discussed in section III-B, we can find the exact population statistics of the model, demonstrated in Fig. 2B and 2C. It is observable that similar to the state-dependent case, for the range of parameters we work with, the proposed analytical approximation fits the numerical solution well, and extinction has a very low probability of occurrence.

## V. Discussion

In this contribution, we have used the discrete stochastic logistic model framework to study the optimal harvesting problem. The main difference of this approach with previous works is that the population is taken as discrete and there is a non-zero probability that the population gets extinct. During harvest, each individual has a probability *f* of being caught. Using the previously developed derivative matching closure scheme, we obtained analytical approximations for different harvest strategies.

The harvesting strategies studied were unified as particular cases of a general harvesting rate *H*(*x*) ∝ *x*^*α*^ with *α* ≥ 0 the control strength and *x* being the population. *α* = 0 defines the constant harvest rate strategy, *α* = 1 is the state-dependent harvest rate where the harvesting rate is proportional to *x* and *α* → ∞ is the threshold-based strategy where harvesting is made when *x* reaches a threshold. Fig. 3 compares the highest achievable mean yield for the different types of harvesting and reveals that the threshold-based strategy is optimal.

**Fig. 3:**
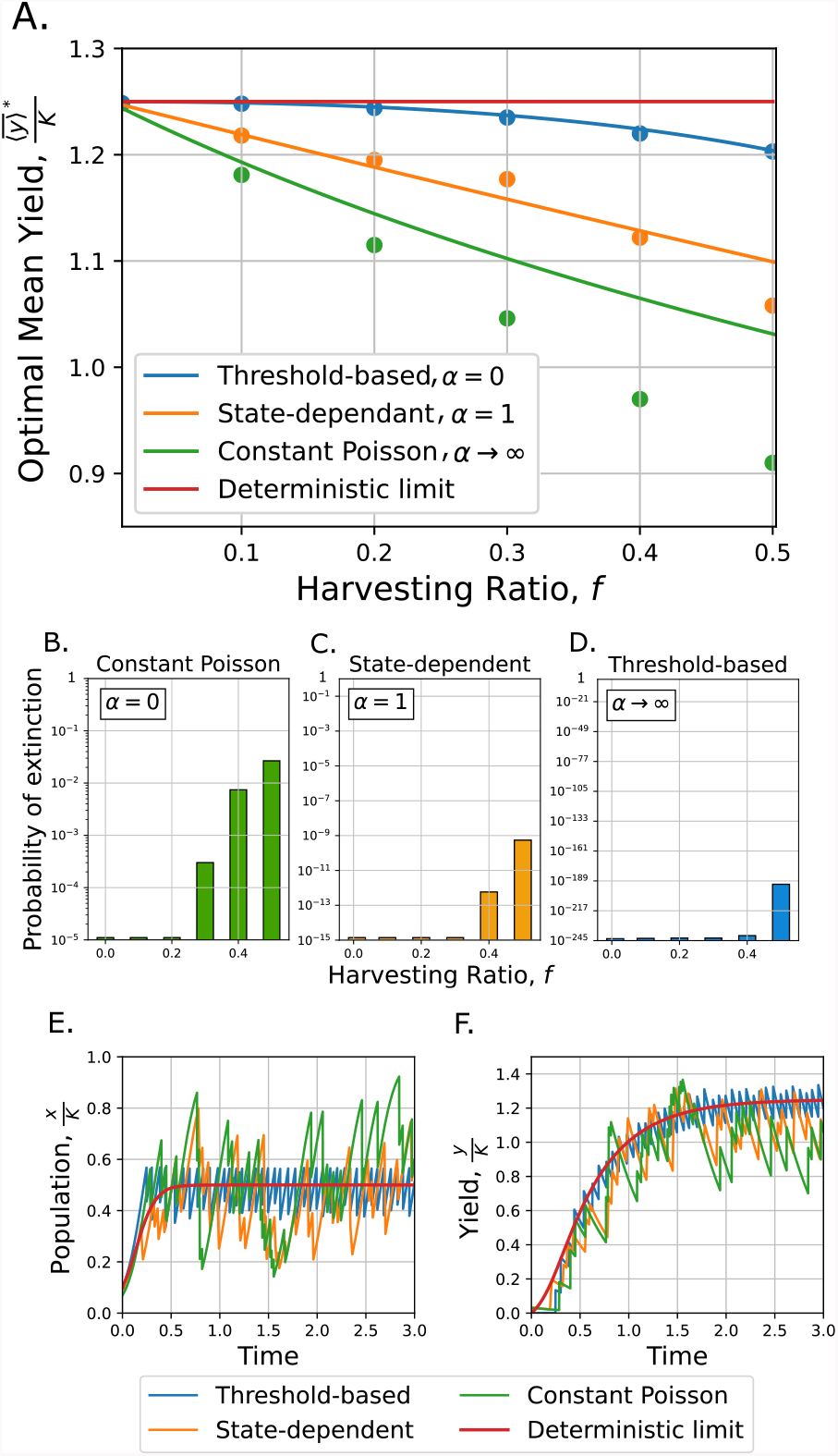
Threshold-based harvesting leads to the maximum yield with lowest probability of extinction comparing to the other two harvesting strategies. **A**. Normalized optimal mean yield 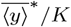 as a function of harvesting ratio *f* for three different harvesting strategies are plotted using the analytic formulas obtained in (12), (19), (26). The dots show the numerical solution by using FSP method for *t*_*max*_ = 400*/r*. The red line shows the normalized deterministic limit of the maximum yield, 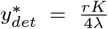 as explained in Section II. Other parameters are taken as *r* = 10, *K* = 10, *λ* = 2. **B, C, D**. The probability of extinction for constant Poisson case increases with the growth of the harvesting ratio *f*, while this probability in state-dependent and threshold-based cases is approximately zero for the given range of parameters. **E, F**. An example of trajectories of population and yield for three harvesting strategies, along with the deterministic limit (*f* = 0.3).

In Fig. 3A, we compare the analytical approximations to the optimal yield 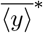 (solid line) and the more accurate numerical solution of the associated Master equation (dots). To study how much of their differences may be due to extinction effects, we present the probability of extinction for the three strategies in Fig. 3B, C, D. For the constant Poisson rate, the probability of extinction can be as high as 0.03 during the studied time scale (400*/r*), with *r* the proliferation rate. For the state-dependent and threshold-based strategies, the probability of extinction is practically null, and the differences between the formula and the numerical result can be due to the approximations made.

We also observe that at the limit of perturbative harvest (*f* → 0), the optimal yield of all strategies converges to the global maximum yield 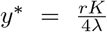 with *K* being the carrying capacity, *r* the exponential proliferation rate, and *λ* the exponential rate of consumption. The value of this yield corresponds to the deterministic limit *y*_*det*_ in (5). In Fig. 3E, F, we present examples of population trajectories for the three harvesting strategies along with the deterministic limit.

The yield is optimized in the deterministic limit when the harvest is controlled such that the population is set to 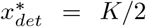. The reason for this value is that the population recovery rate (the rate at which the population grows) is maximized when *x* has this value. When harvesting is modeled by non-perturbative jumps *f* > 0, the threshold-based strategy allows a maintenance of the population around *x*\* better than the state-dependent strategy, resulting in a higher optimal yield. The constant Poisson strategy, on the other hand, does not have population information to decide when to harvest, leading to a relatively higher probability of population extinction in most parameter regimes.

## VI. Appendix

### A. Estimation of the formulas of the statistics for the threshold based strategy

In the regime of a high population number, *K* ≫ 1 we approximate the dynamics of the population using the logistic curve:

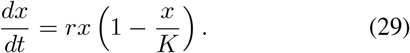

Once the population reaches the threshold *x*_*h*_, the population is reset to a new variable *x*_0_. In order to mimic the discrete stochastic harvesting, we assume that *x*_0_ follows a statistic similar to a binomially distributed variable:

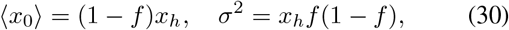

with *σ*^2^ being the variance. Given *x*_0_, the dynamics of the population follows the solution:

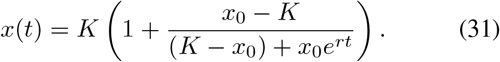

Hence, the time *τ* (*x*_0_) for the next harvest is the time needed to reach *x*_*h*_ again:

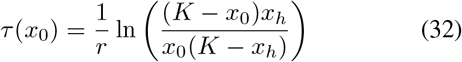

Once *x*_0_ is a random variable, it is possible to show that the average of a multiple harvests can be written as [36]

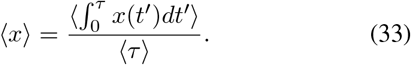

with ⟨*f* ⟩ = ∫ *ρ*(*x*_0_)*f* (*x*_0_)*dx*_0_ the expected value of a function *f* (*x*_0_) over the distribution *ρ*(*x*_0_) of *x*_0_. Given the moments of *x*_0_, we can simplify the averaging considering *x*_0_ drawn from a uniform distribution with statistical moments (30).

To simplify the estimation, we assume that *σ << K*. This, considering (30), is equivalent to considering *x*_*h*_ << *K*^2^. Therefore, the average (33) can be expanded in power series:

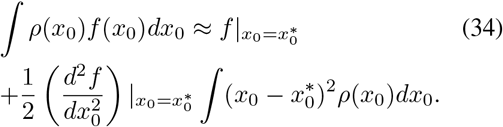

Where the integration of the first order of the polynomial expansion is null since the distribution is odd around 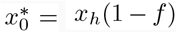. After these approximations, the mean population can be approximated as:

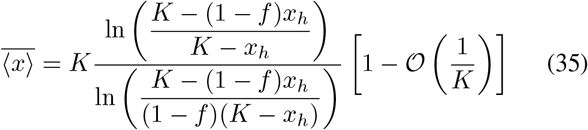

with:

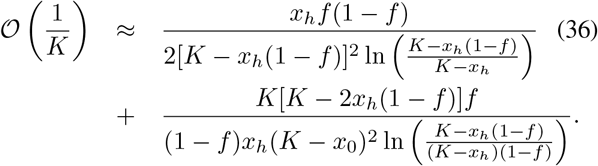

